# Trem2 Agonist Reprograms Foamy Macrophages to Promote Atherosclerotic Plaque Stability

**DOI:** 10.1101/2023.09.21.558810

**Authors:** Michael T. Patterson, Yingzheng Xu, Hannah Hillman, Victoria Osinski, Patricia R. Schrank, Ainsley E. Kennedy, Alisha Zhu, Samuel Tollison, Sia Shekhar, Ingunn M. Stromnes, Ilaria Tassi, Dick Wu, Bryce A. Binstadt, Jesse W. Williams

## Abstract

**Objective:** Trem2, a surface lipid receptor, is expressed on foamy macrophages within atherosclerotic lesions and regulates cell survival, proliferation, and anti-inflammatory responses. Studies examining the role of Trem2 in atherosclerosis have shown that deletion of Trem2 leads to impaired foamy macrophage lipid uptake, proliferation, survival, and cholesterol efflux. Thus, we tested the hypothesis that administration of a validated Trem2 agonist antibody (AL002a) to atherogenic mice could drive macrophage survival and decrease necrotic core formation to improve plaque stability.

**Approach and Results:** To model a therapeutic intervention approach, atherosclerosis-prone mice (Ldlr^-/-^) were fed a high fat diet (HFD) for 8 weeks, then transitioned to treatment with AL002a or isotype control for an additional 8 weeks while continuing on an HFD. AL002a-treated mice had increased lesion size in both the aortic sinus and whole mount aorta, which correlated with an expansion of plaque macrophage area. This expansion was due to increased macrophage survival and proliferation in plaques. Importantly, plaques from AL002a-treated mice showed improved features of plaque stability, including smaller necrotic cores, increased fibrous caps, and greater collagen deposition. Single cell RNA sequencing of whole aorta suspensions from isotype and AL002a-treated atherosclerotic mice revealed that Trem2 agonism dramatically altered foamy macrophage transcriptome. This included upregulation of oxidative phosphorylation and increased expression of collagen genes. In vitro studies validated that Trem2-agonism with AL002a promoted foamy macrophage oxLDL uptake, survival, and cholesterol efflux in culture.

**Conclusions:** Trem2 agonist expands plaque macrophages by promoting cell survival and proliferation but improves features of plaque stability by rewiring foamy macrophage function to enhance collagen deposition.

## Introduction

Atherosclerosis is a chronic inflammatory disease of the arteries that is a major cause of cardiovascular disease. Atherosclerotic plaque progression is driven by the deposition of pathogenic low-density lipoprotein (LDL) within the intima that is internalized by plaque macrophages, forming lipid-engorged foamy cells^1^. Foamy macrophages are a hallmark of atherosclerotic plaques and are one of the most abundant cell types present within early lesions. Importantly, accumulation of excess LDL within foamy macrophages is cytotoxic^2–4^, driving apoptosis and ultimately formation of necrotic cores. Necrotic cores typically consist of masses of dead cells along with extracellular lipid and are a result of excess macrophage apoptosis^5,6^ and impaired dead cell clearance (efferocytosis)^7,8^. Advanced atherosclerotic plaques are characterized by having large necrotic cores^9,10^ and this, along with impaired fibrous cap development^11^, contributes to unstable lesions that are prone to rupture and cause myocardial infarction and stroke^12,13^.

Recent efforts targeting macrophages to improve features of plaque stability are underway. Studies in preclinical models and humans have definitively shown that increasing macrophage efferocytosis^14–16^ and promoting cell survival, particularly through driving Liver X Receptor (LXR) mediated cholesterol efflux to HDL^17–19^, leads to decreased necrotic core formation and improved features of plaque stability. Furthermore, rewiring macrophage phenotypes toward an alternatively activated state to promote anti-inflammatory cytokine production and extracellular matrix remodeling has been effective in driving plaque regression^20^.

Interestingly, transcriptional analysis of single cell RNA-sequencing (scRNA-seq) data revealed that foamy macrophages within atherosclerotic plaques express the phagocytic receptor Trem2^21^. Trem2 is a surface lipid receptor primarily expressed on tissue macrophages and microglia and has been proposed to activate Syk, PI3K, AKT, and mTOR pathways to drive cell survival, proliferation, efferocytosis and anti-inflammatory function^22–24^. Trem2 has been extensively studied in Alzheimer’s disease models due to its role in promoting microglia survival^22^, and Trem2 agonist antibodies have been shown to be efficacious in slowing disease progression^25^. However, recent works interrogating the role of Trem2 in atherosclerosis in murine models have suggested context dependent results. Macrophage specific deletion of Trem2 during early plaque development slowed plaque progression due to impaired foamy macrophage proliferation and survival through decreased LXR-mediated cholesterol efflux^26^. However, in advanced plaques, whole body Trem2 deletion led to increased necrotic core formation due to impaired efferocytosis and excess cell death^27^. Overall, these data suggest that while inhibiting Trem2 during early lesion development may slow disease progression, promoting Trem2 signaling in advanced disease could impair necrotic core formation and plaque inflammation to promote stability.

Here, using a therapeutically relevant Trem2 agonist (AL002a, a murine surrogate antibody for Trem2)^25^, we interrogated the effects of promoting Trem2 signaling in a model of atherosclerosis. We find that treatment with AL002a promoted plaque macrophage expansion, proliferation, and survival while simultaneously decreasing necrotic core formation and improving fibrous cap development. Using scRNA-seq transcript analysis of atherosclerotic aortae and in vitro studies, we determined that Trem2 agonism dramatically rewired foamy macrophage metabolism, promoted foamy cell collagen production, and drove cholesterol efflux to improve features of plaque stability.

## Materials and Methods

### Study Design

In vivo studies of atherosclerosis used both male and female Ldlr^-/-^ mice (n=7-10/group) (B6.129 S7-Ldlr^tm1Her^/J, Jax 002207) aged to 7 weeks prior to enrollment. At 7 weeks of age, littermate mice were started on a high fat diet (HFD) (diet no. TD.88137; adjusted calories diet, 42% from fat, Envigo Teklad) for 8 weeks then randomized to receive either Isotype control or Trem2 agonist at 40 mg/kg/week (AL002a, Alector Inc.^25^) i.p. weekly for an additional 8 weeks. A terminal bleed was performed after 16 weeks on an HFD and was used for serum lipid quantification and flow cytometry. All mice were bred in specific pathogen-free animal facilities maintained by the University of Minnesota (UMN) Research Animal Resources (RAR). All experiments were approved by the UMN Institutional Animal Care and Use Committee (IACUC).

### Atherosclerosis and Necrotic Core Analysis

To examine atherosclerotic plaque size, aortae and hearts were harvested from mice and fixed in 4% PFA overnight. Hearts were embedded in OCT, frozen then sectioned on a cryostat at 10 μm thickness. Sections were stained with Oil Red-O and hematoxylin, and images captured using Leica SP8 inverted confocal microscope. For aortae, after fixation, samples were cut open and pinned en face to wax dishes. Samples were washed with water, then put in propylene glycol for 5 minutes. Next, the aortae were incubated in Oil Red O (Sigma O1516) for three hours. Afterwards the dishes were washed in 85% propylene glycol for 5 minutes followed by water. Images were taken using Leica S9i stereo microscope with 10 megapixel camera. Lesion area was quantified using ImageJ and blinded-analysis on 7 serial sections from each mouse. Necrotic core analysis was quantified by blinded investigators using ImageJ. Necrotic core area was defined as acellular areas within plaque sections as previously published^16^.

### Serum Cholesterol Analysis

Blood was clotted at room temperature for 1 hour, then samples were centrifuged. The supernatant (serum) was collected and assessed for cholesterol content using Wako/Fujifilm Cholesterol-E kit (#999-02601), per manufacturer’s instructions.

### low Cytometry

Whole blood was lysed, and single cell suspensions were filtered through 100 μm nylon mesh, then washed in FACS buffer (HBSS with 2% FBS and 2mM EDTA). Cells were then stained for 30 minutes at 4°C, protected from light. Antibodies were stained at 1 mg/mL in 100 μL. Spectral cytometry was collected using a Cytek Aurora. All machines are supported and maintained by the UMN flow cytometry core facility. Data was assessed in Flowjo (Tree Star). The following antibodies were used: Anti mouse CD45 BV480 (Clone: 30-F11, BD, Cat#:566168), Anti mouse CD11b BV605 (Clone: M1/70, Biolegend, Cat#:101237), Anti mouse Ly6G BV785 (Clone: 1A8, Biolegend, Cat#:127645), Anti mouse Ly6C BV421 (Clone: HK1.4, Biolegend, Cat#:128031), Anti mouse CD115 PerCp Cy5.5 (Clone: AFS98, Biolegend, Cat#:135525), Anti mouse TCRβ APC (Clone: H57-597, Biolegend, Cat#:109211) and Anti mouse CD19 FITC (Clone: ID3, Biolegend, Cat#:152403).

### Immunostaining of Aortic Sinus

Slides were warmed to room temperature for 10 minutes on the bench top. Samples were washed with PBS then blocked with 5% donkey serum and permeabilized with 1% triton x-100 for 20 minutes. Primary antibodies were diluted 1:500 in PBS and samples were stained for 1 hour in dark. Samples were washed three times with PBS, then stained with secondary conjugated antibodies (1:500 dilution) for 1 hour. Samples were washed 3x, then mounted with flouromount (Southern Biotec). Samples were imaged using a Leica SP8 inverted confocal microscope (fluorescence imaging). For macrophage identification and area quantification, CD68 (Clone: FA-11, Biolegend, Cat#:137001) was used. Area was determined using ImageJ. For Tunel staining, a Roche In Situ Cell Death Detection Kit (Millipore Sigma) according to the manufacturer’s instruction was used and CD68+ Tunel+ cells were counted. For proliferation, anti-mouse Ki67 (Clone: SP6, Abcam, Cat#: ab16667) was used and CD68+ Ki67+ cells were counted. SMA staining was performed using anti mouse SMA and area of SMA within fibrous cap was quantified using ImageJ as previously published^28^. For picrosirius red staining, slides were warmed to room temperature for 10 minutes on the bench top. Sections were then hydrated in distilled water and stained with Sirius red (0.1% Direct Red 80 (Sigma, 365548-5G) in saturated aqueous solution of picric acid (Ricca Chemical, 5860-16)) for one hour at room temperature. Slides were then dipped in 0.5% acetic acid, dehydrated in 95% ethanol then 100% ethanol, and finally cleared in xylene. Images were captured using polarized light on a Stellaris 8 microscope (Leica). Using 8-bit images, regions of interest were drawn around each plaque in ImageJ. Following thresholding, the percent of positive collagen signal was quantified per plaque.

### Whole Aorta scRNAseq

scRNAseq was performed on digested, whole aorta from either isotype or AL002a treated mice after 16 weeks HFD (n=7/group). Both male and female mice were used, with overlapping phenotypes observed. Briefly, aortae were digested and single cell suspensions were generated as previously published^29^. Samples were then stained with hashtag oligo antibodies (BD Biosciences; HTO#9 and HTO#10) and then resuspended in a final concentration of 100 cells per μl in PBS for single cell capture of approximately 20,000 cells per group. Cells were submitted to University of Minnesota Genomics Core (UMGC) for single cell 10X Chromium 3’ GEX Capture and NovaSeq 2 x150 S4 sequencing targeting ∼50,000 reads per cell.

Preparation and generation of count matrices was provided by the UMGC using Cell Ranger. Captures were integrated using the Harmony package (v0.1.0). Further downstream analysis was performed using the Seurat package (v4.2.1). Genes expressed by less than 10 cells were removed. Ribosomal genes were excluded from further analysis as they heavily skew differential expression (DE) analysis. Cells of inferior quality were determined and filtered based on two criteria: total read counts below 300, and fraction of mitochondrial genes greater than 25%. Doublets were identified using the DoubletFinder package (v2.0.3). Post principal component analysis by RunPCA, RunTSNE function was employed to project cells in a reduced dimensional space. Clusters were finetuned using FindNeighbors and FindClusters with resolution 0.2. The myeloid subset was then isolated for reclustering using resolution 0.3. DE analysis was carried out using FindMarkers or FindAllMarkers. Log-fold change ranked genes were used as background for pathway analysis through the fgsea package (v1.18.0). In this context, the Reactome or Hallmark pathway libraries were primarily utilized. Furthermore, data visualization was facilitated by the ggplot2 package (v3.4.0).

### BMDM Generation

Mouse bone marrow was isolated from the femur and tibia of C57BL/6 mice, prepared as a single-cell suspension, and red blood cells lysed for 1 minute using ACK lysis buffer. Cells were plated in CMG conditioned media: DMEM (Gibco) with 10% FBS, 1% PenStrep (Sigma-Aldrich), 1% Sodium Pyruvate (Millipore), 1% L-glutamine (Sigma), 10% CMG MCSF cell supernatant.

Cells were maintained in a 37ºC incubator with 5% CO2. Media was refreshed on Day 4. On Day 7, cells were removed from plate with trypsin-EDTA (Gibco) and used in in vitro assays. Macrophage identity was confirmed via flow cytometry by confirming expression of F4/80 (BV650, Biolegend, Cat#: 123149) and CD11b (APC Fire 750, Biolegend, Cat#: 101262).

### Cholesterol Efflux Assay

BMDMs were aliquoted at 200,000 cells/ml and treated with either isotype control, LXR agonist (T0901317, 10μM), or AL002a (2nM). All cells received soluble cholesterol at 20 μg/ml (Millipore, Cat#: C4951-30MG) to differentiate into foamy macrophages, and then aliquoted into a 24-well plate (1ml mixture/well). Cells were incubated for 16 hours overnight in a 37ºC incubator with 5% CO_2_. The following day, cells were scraped from the wells, washed in HBSS (Sigma-Aldrich), and stained with BODIPY lipid dye at 1:1000 in HBSS for 15 minutes at 4ºC. Cells were washed, 10% FBS was added to act as a cholesterol acceptor, and cells effluxed for 4 hours at 4ºC. Cells were then acquired on the BD LSRII flow cytometer and data analyzed using FlowJo.

### LDH Release Assay

Cell viability was assessed using the Cytotoxicity Detection Kit Plus (LDH) (Roche, Cat#: 04744926001) according to manufacturer’s instructions. Briefly, BMDMs were incubated overnight in a 96-well plate (50k cells/well) in media alone or with 20 μg/ml soluble cholesterol, and with either isotype control or AL002a. The following day, prepared reaction mixture was added to the cells for 20 minutes at room temperature. Positive controls also received 5 μl lysis buffer for an additional 15 minutes. All wells were then treated with 50 μl stop solution, and absorbance at 490nm was measured using the TECAN Infinite 200 Pro microplate reader. Percent cytotoxicity was determined using the absorbance values minus the background controls and normalized to baseline per the manufacturer:

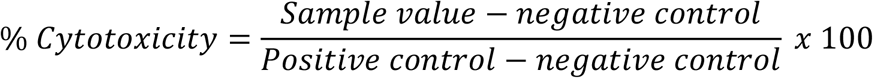

### oxLDL Uptake Assay

BMDMs were incubated overnight in a 24-well plate in media alone or with 20 μg/ml soluble cholesterol, then treated with either isotype control or AL002a. The following day, DiI-oxLDL (Kalen Biomedical, Cat#: #770282-9) was added at 10 μg/ml for 4 hours at 37ºC. Cells were lifted using trypsin-EDTA, washed in HBSS, and stained for flow cytometry analysis.

#### Statistical Analysis

Graphs were generated and statistical analysis performed in Graphpad Prism. For a comparison between two experimental groups a student’s t-test was used, whereas comparisons of more than 2 groups utilized a two-tailed ANOVA. Graph error bars represent standard error of the mean (SEM), and *P*-value was statistically significant below <0.05. In graphs, *<0.05, **<0.01, ***<0.001, ****<0.0001.

## Results

### Trem2 agonist AL002a expands plaque macrophages but improves features of stability

Recent reports indicate that Trem2 deficiency impairs foamy macrophage survival through impaired cholesterol efflux^26^ and exacerbates necrotic core formation^27^. Thus, we hypothesized that promoting Trem2 signaling in atherosclerosis could drive macrophage survival, decrease plaque necrotic core formation, and promote stable plaque phenotypes. To test this hypothesis, we treated atherogenic Ldlr^-/-^ mice with a previously validated Trem2 agonist antibody AL002a, a surrogate mouse antibody for Trem2^25,30^. In a therapeutic model for treating established atherosclerosis, mice were fed an HFD for 8 weeks, then treated with either isotype control or AL002a for an additional 8 weeks while continuing HFD feeding (Figure 1A). After 16 weeks of total HFD, mice were sacrificed and assessed for atherosclerosis progression via Oil Red O (ORO) staining of whole mount aortae and aortic sinus cross sections. Aortic sinus analysis revealed that treatment with AL002a lead to an expansion of plaque sizes regardless of distance through the root (Figure 1B-C). Furthermore, whole mount aorta ORO staining confirmed that AL002a treatment increased the total atherosclerotic plaque burden (Figure 1D-E). Importantly, AL002a treatment of HFD fed mice did not alter body weight accumulation, cholesterol levels or blood immune cell proportions compared to isotype control treatment (Supplemental Figure 1).

**Figure 1.**
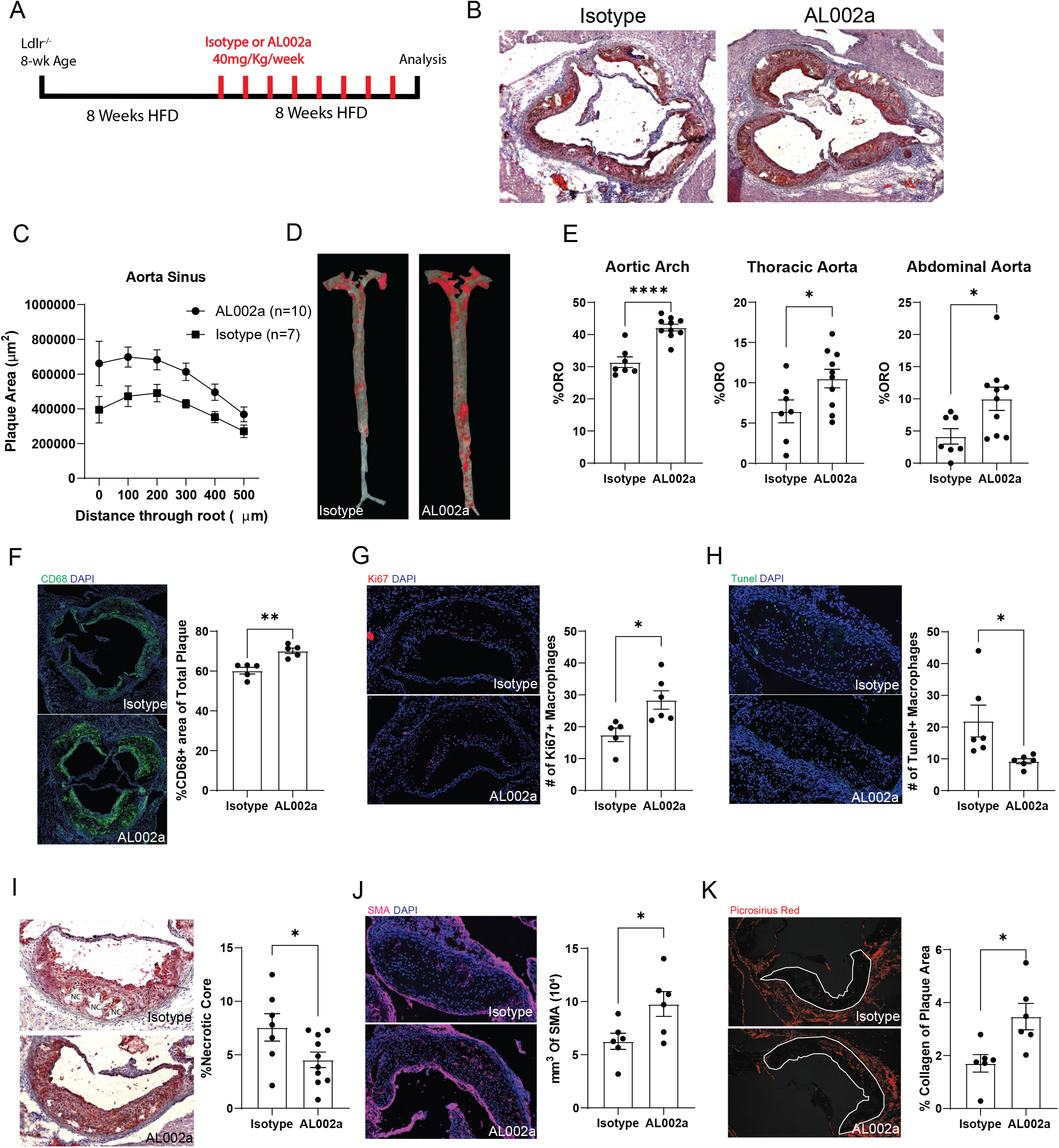
Trem2 agonist AL002a expands plaque macrophages but improves features of stability. A) Scheme of experimental approach. Ldlr^-/-^ mice were fed a high fat diet (HFD) for 8 weeks then treated with either isotype control or Trem2 agonist (AL002a) weekly for an additional 8 weeks of HFD at 40 mg/kg. B) Representative images of aortic root cross sections stained with oil red o (ORO) from isotype and AL002a treated mice after 16 weeks HFD. C) Aortic root area measurement by distance through root after ORO staining from isotype and AL002a treated mice (n=7-10/group). Data are mean ± S.E.M. D) Representative en face whole aorta stained with ORO from isotype and AL002a treated mice after 16 weeks HFD. E) Arch, thoracic, and abdominal aorta analyzed by en face analysis for percentage oil red O (ORO) staining after 16 weeks HFD (n=7-10/group). Data are mean ± S.E.M. F) Representative images and quantification of CD68+ macrophage area of aortic root cross sections after 16 weeks HFD from isotype and AL002a treated mice (n=5/group). CD68 in green, DAPI in blue. Data are mean ± S.E.M. G) Representative images and quantification of Ki67+ macrophages in aortic root cross section plaques after 16 weeks HFD from isotype and AL002a treated mice (n=5-6/group). Ki67 in red, DAPI in blue. Data are mean ± S.E.M. H) Representative images and quantification of TUNEL+ macrophages in aortic root cross section plaques after 16 weeks HFD (n=7/group). TUNEL in green, DAPI in blue. Data are mean ± S.E.M. I) Necrotic core quantification and images in aortic root cross section plaques after 16 weeks HFD (n=7-10/group). Necrotic cores were identified using acellular areas from hematoxylin and ORO-stained sections. Data are mean ± S.E.M. J) Smooth muscle fibrous cap area quantification and images from aortic cross sections after 16 weeks HFD (n=6/group) using smooth muscle actin (SMA). SMA in pink, DAPI in blue. Data are mean ± S.E.M. K) Picrosirius red polarized light images and percentage quantification for collagen content from aortic cross sections after 16 weeks HFD (n=6/group). Plaque area outlined and picrosirius red in red. Data are mean ± S.E.M.

Given that Trem2 deletion leads to decreased macrophage cellularity within atherosclerotic plaques through impaired macrophage survival and proliferation^26^, we next assessed plaque macrophage phenotype. CD68 staining of aortic root cross sections of HFD fed mice revealed that AL002a treatment expanded macrophage area within plaques compared to isotype control treatment (Figure 1F). To determine mechanisms driving the expansion of macrophages within plaques of AL002a treated mice, macrophage proliferation and survival was examined by confocal microscopy. Ki67 immunostaining of aortic root cross sections of HFD fed mice found that AL002a treatment led to an increase in the number of proliferating macrophages (Ki67+ CD68+) within plaques (Figure 1G). TUNEL staining was performed to identify apoptotic cells and found that AL002a treated mice had a dramatic reduction in the number of dying plaque macrophages (Figure 1H), suggesting that Trem2 agonism promotes macrophage expansion within plaques by driving their proliferation and persistence within lesions.

Next, we sought to examine features of plaque stability by measuring necrotic core and fibrous cap formation from atherosclerotic mice treated with isotype control or AL002a. Analysis of necrotic core within aortic root plaques revealed that AL002a treatment led to an ∼25% decrease in necrotic core area compared to isotype controls (Figure 1I). This is consistent with AL002a’s ability to promote macrophage survival. Smooth muscle fibrous cap formation was also quantified as a readout of plaque stability by staining aortic root cross sections for smooth muscle actin (SMA). SMA cap area was expanded following Trem2-agonist treatment (Figure 1J). Consistent with this, collagen deposition within lesions was also measured using picrosirius red, where plaques from AL002a treated mice had an increase in collagen content compared to isotype control treated mice (Figure 1K). Overall, these data suggest that AL002a treatment expands plaque macrophages survival and improves fundamental features of plaque stability including decreased necrotic core formation and increased fibrosis.

### AL002a reprograms foamy macrophages in atherosclerotic plaques

To determine how AL002a changes plaque cellular phenotypes, we performed scRNAseq on single cell preparations of whole aortas from Ldlr^-/-^ mice fed HFD for 16 weeks and treated with isotype control or AL002a for 8 weeks (Figure 2A). Raw data were integrated, normalized, and clustered using Seurat and cell types were identified using SingleR^31^. These data revealed a diversity of both immune and non-immune cells residing within atherosclerotic aorta, including stomal cells, fibroblasts, myeloid and lymphoid cells (Figure 2B, Supplemental Figure 2A).

**Figure 2.**
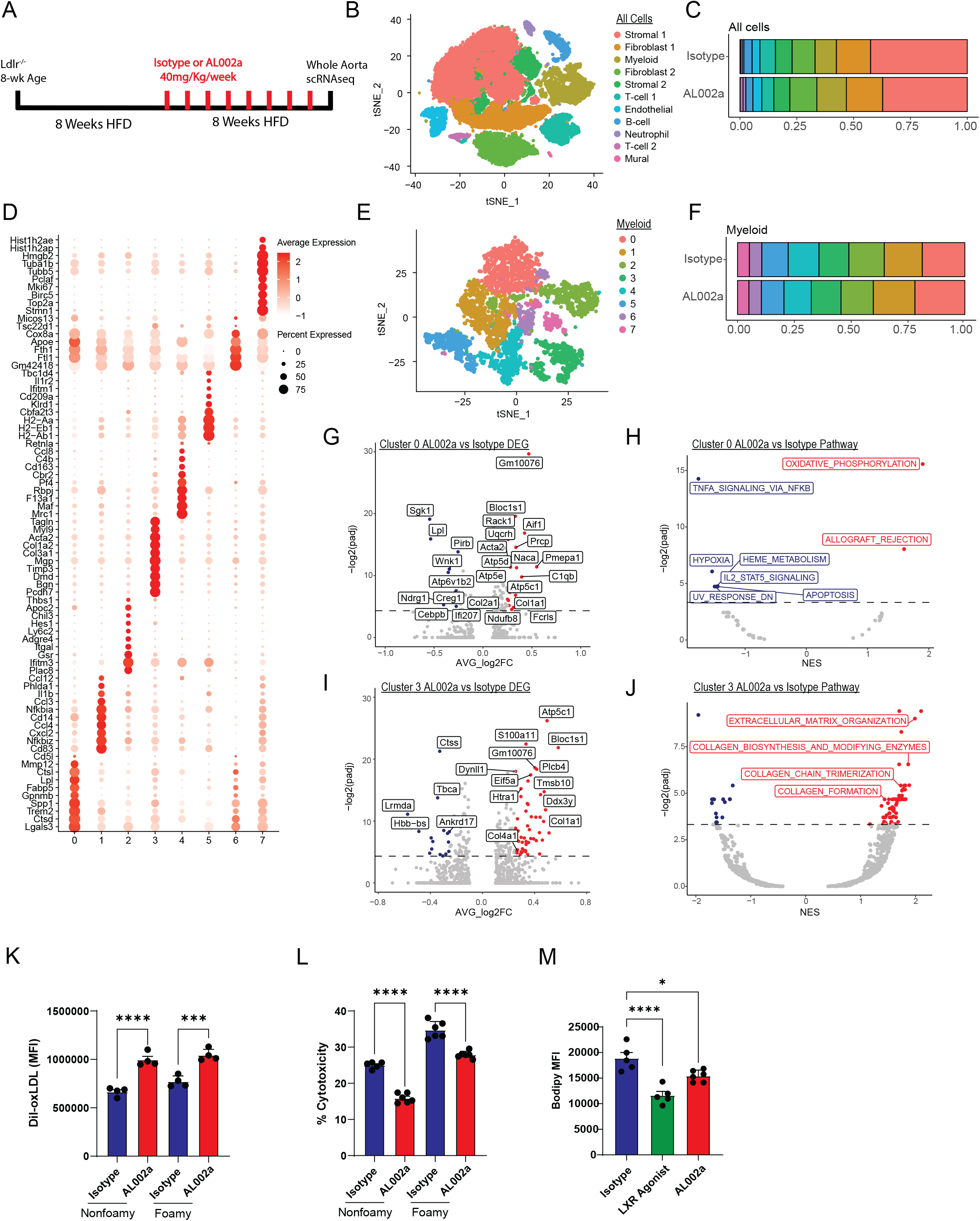
AL002a reprograms foamy macrophages in atherosclerotic plaques. A) Scheme of experimental approach. Ldlr^-/-^ mice were fed a high fat diet (HFD) for 8 weeks then treated with either isotype control or Trem2 agonist (AL002a) weekly for an additional 8 weeks of HFD. Mice were sacrificed at 8 weeks and aortae were harvested, digested and sequenced (n=7/group). B) tSNE clustering of total cells from atherosclerotic aortae from both isotype and AL002a treated mice (∼30,000 cells). Cell clusters were named using SingleR. C) Proportion of each cluster from B split by treatment group. D) Heatmap showing normalized cluster-specific gene expression from re-clustering of myeloid population in B. E) tSNE clustering of only myeloid cells from aortae from both isotype and AL002a treated mice showing 8 unique myeloid subpopulations. F) Proportion of each myeloid cluster from E split by treatment group. G) Volcano plot showing differentially expressed genes between foamy macrophage cluster (cluster 0) of isotype and AL002a treated mice. Genes to the right (red dots) are upregulated in AL002a foamy macrophages, genes to the left (blue dots) are upregulated in isotype control foamy macrophages. H) REACTOME pathway analysis of differentially expressed genes from G. Pathways to the right (red dots) enriched in AL002a foamy macrophages, pathways to the left (blue dots) are enriched in isotype control foamy macrophages. I) Volcano plot showing differentially expressed genes between smooth muscle foamy cell cluster (cluster 3) of isotype and AL002a treated mice. Genes to the right (red dots) are upregulated in AL002a smooth muscle foamy cells, genes to the left (blue dots) are upregulated in isotype control smooth muscle foamy cells. J) REACTOME pathway analysis of differentially expressed genes from I. Pathways to the right (red dots) enriched in AL002a smooth muscle foamy cells, pathways to the left (blue dots) are enriched in isotype control smooth muscle foamy cells. K) Non-foamy and foamy bone marrow derived macrophages (BMDMs) were pretreated with either isotype control or AL002a (0.85 ul/mL) in culture for 4 hours then DiI-oxLDL was added to the media for an additional 4 hours (n=4/group). Uptake was then quantified by flow cytometry. Data are mean ± S.E.M. L) BMDMs were incubated overnight in media alone (non-foamy) or with 20 mg/mL of soluble cholesterol (foamy) and with either isotype or AL002a antibody (n=5/group). Supernatants were then harvested and ran for lactate dehydrogenase (LDH) concentration as a readout of cytotoxicity. Data are mean ± S.E.M. M) BMDMs were treated with 20 mg/mL of soluble cholesterol overnight, stained with BODIPY to label intracellular cholesterol then re-cultured in serum free media plus HDL cholesterol acceptor with either isotype antibody, LXR agonist T0901317 or AL002a for 4 hours for effluxing. Cells were measured for loss of BODIPY for efflux ability (n=4-5/group). Data are mean ± S.E.M.

Differential gene analysis of cluster defining genes shows that both stromal cell clusters express high levels of *Acta2* and *Myl9*, suggesting that these populations are smooth muscle cell derived (Supplemental Figure 2A). Furthermore, the myeloid cell cluster highly expressed genes associated with macrophage phenotype (*Lyz2, Apoe*) implying that this cluster primarily consists of macrophages (Supplemental Figure 2A). Next, we compared the proportion of each cell population between isotype and AL002a treated mice and found that AL002a treatment increased the proportion of myeloid cells compared to isotype control treatment (Figure 2C), consistent with Trem2 agonism driving macrophage expansion.

To compare changes of macrophage subpopulations, we subclustered myeloid cells from isotype and AL002a treated aortas and found 8 distinct populations (Figure 2D-E). Similar to prior meta-analyses of scRNAseq studies in atherosclerosis^21^, we identified a foamy macrophage population (Cluster 0) that expressed genes associated with lipid loading (*Lgals3, Fabp5, Trem2*), an inflammatory macrophage population (Cluster 1) that expressed proinflammatory genes (*Il1b, Cd83, Cxcl2*) and a monocyte population (Cluster 2) that expressed canonical monocyte genes (*Ly6c2, Plac8*) (Figure 2D). Interestingly, cluster 3 co-expressed genes associated smooth muscle cells (*Acta2, Myl9*) and foamy cells (*Lgals3, Fabp5*), which is consistent with a smooth muscle derived foamy cell phenotype^32^ (Figure 2D, Supplemental Figure 2B). Cluster 4 was enriched for genes associated with adventitia resident macrophages (*Mrc1, Cd163*), while cluster 6 expressed genes of mixed origins suggesting a transitioning macrophage population (Figure 2D). Furthermore, cluster 8 largely consisted of proliferating macrophages due to their high expression of cell cycle and division genes (*Top2a, Mki67*). Cluster 5 expressed high levels of MHC class II genes (*H2-Aa, H2-Ab1*) and *Cd209a* delineating this population as dendritic cells. Comparing proportions of these populations between isotype control and AL002a treated mice revealed relatively minor differences between treatments, with AL002a treatment leading to an expansion of foamy macrophages (Figure 2F).

Next, we sought to examine gene expression differences driven by AL002a treatment between macrophage subpopulations. We hypothesized that Trem2-expressing cells would be the most affected by AL002a treatment and found that foamy macrophages expressed the highest level of Trem2 (Supplemental Figure 2B). Surprisingly, we also found that smooth muscle derived foamy cells expressed considerable levels of Trem2 (Supplemental Figure 2B), albeit lower than monocyte-derived foamy cells, suggesting that they could also be reprogrammed by AL002a.

Differential gene analysis of foamy macrophages (Cluster 0) from isotype- and AL002a-treated mice revealed dramatic alterations in genes associated with lysosomal function and cell metabolism, with AL002a-treated foamy macrophages showing increased expression of lysosomal (*Bloc1s1, Prcp*) and oxidative phosphorylation genes (*Atp5c1, Atp5e, Uqcrh*).

Furthermore, foamy macrophages from AL002a-treated plaques expressed higher levels of genes involved in collagen production (*Col1a1, Col2a1*) compared to isotype control-treated mice. Pathway analysis of differentially expressed genes from foamy macrophages revealed that AL002a treatment led to an enrichment in oxidative phosphorylation and a downregulation in pathways involved in inflammation and apoptosis (Figure 2H), suggesting that Trem2 agonism rewires foamy macrophage metabolism, promotes cell survival, and dampens inflammation. We also assessed differentially expressed genes between smooth muscle derived foamy cells (Cluster 3) and found that AL002a treatment again led to increased expression of genes involved in collagen production (*Col1a1, Col4a1*) (Figure 2I). This was also reflected by an enrichment in pathways involved in ECM organization and collagen production following AL002a treatment (Figure 2J, Supplemental Figure 2C). Overall, this analysis supports that Trem2 agonism dramatically rewires both monocyte- and smooth muscle-derived foamy macrophage function in lesions to promote features of plaque stability including cell survival, modulation of inflammation, and improved ECM production.

To elucidate gene changes in other cell subsets with Trem2 agonism, we also assessed differentially expressed genes in stromal and fibroblast clusters between plaques from isotype and AL002a treated mice. Since these cell types do not express Trem2, it is expected that effects observed in these cell types are a downstream consequence of reprograming foamy macrophages within the local environment. Differential gene and pathway analysis found several alterations, most notably decreased fibroblast proliferation and increased smooth muscle collagen production (Supplemental Figure 3), supporting the notion that driving Trem2 signaling in Trem2+ cells can reshape other artery cell subsets.

Lastly, to examine how Trem2 agonism modifies macrophage function, we performed in vitro assays in bone marrow derived macrophages (BMDMs) treated with either isotype control or AL002a antibodies. BMDMs were differentiated with M-CSF then treated overnight in media alone or 20 μg/mL soluble cholesterol to generate nonfoamy or foamy macrophages, respectively, and treated with 2nM Trem2 agonist AL002a or isotype control antibody. First, BMDMs treated with or without AL002a were assessed for ability to uptake oxLDL by treating cells with fluorescently labeled DiI-oxLDL for 4 hours. AL002a treatment led to an increased ability of macrophages to take up oxLDL in both the non-foamy and foamy state (Figure 2K), which is supported by the expansion in foamy macrophages observed in vivo (Figure 2F). Next, given the proposed role of Trem2 in promoting myeloid cell survival under chronic phagocytic challenge by driving cholesterol efflux^26,33^, we assessed cytotoxicity and cholesterol efflux in vitro. Pretreatment of BMDMs with AL002a was sufficient to increase cell viability in nonfoamy and foamy cells (Figure 2L). To assess cholesterol efflux, BMDMs were cultured with 20 μg/mL soluble cholesterol overnight with either isotype or AL002a. Cells were then stained with the cholesterol dye BODIPY and allowed to efflux for 4 hours in media containing FBS as a cholesterol acceptor. To measure efflux, BODIPY loss was assessed via flow cytometry. As a positive control, BMDMs treated with the Liver X Receptor (LXR) agonist T0901317 were also assessed. Treatment with either an LXR agonist or AL002a decreased BODIPY fluorescence intensity after 4 hours, consistent with an ability to enhance cholesterol efflux (Figure 2M). In conclusion, these in vitro studies complement and support our in vivo studies to conclude that Trem2 agonism promotes key features of foamy macrophage function, including macrophage oxLDL uptake, survival, and cholesterol efflux.

## Discussion

Here, we tested the role of Trem2 agonism on established atherosclerotic lesions. Treatment of atherosclerotic mice with Trem2 agonist rewired foamy macrophages within atherosclerotic plaques which led to increased lesion size through promotion of macrophage survival. However, Trem2 agonist also promoted fundamental features of plaque stability, including increased smooth muscle cap formation, increased collagen deposition, and reduced necrotic core formation. By examining transcriptomic profiles using scRNA-seq, we found that AL002a treatment redirected foamy macrophage metabolism toward oxidative phosphorylation and promoted collagen production. Finally, in vitro studies of foamy and nonfoamy BMDMs revealed that driving Trem2 signaling can promote uptake of oxLDL, cell survival, and cholesterol efflux.

We previously demonstrated that conditional deletion of Trem2 on plaque macrophages led to decreased lesion size, primarily through decreased macrophage proliferation and survival^26^. In line with this, we now demonstrate that Trem2 agonist treatment expands plaque macrophages by promoting macrophage proliferation and persistence within plaques, which we hypothesize is the primary factor leading to plaque expansion. This conclusion is further supported by a recent report investigating germline Trem2 deletion in advanced atherosclerotic plaques that found while Trem2 deletion had no effect on plaque size, Trem2 deficient mice had larger necrotic cores^27^. It has been proposed that the effect of macrophage cell death on plaque outcomes depends on the stage of plaque development^34,35^. Thus, we hypothesize that while promoting foamy macrophage cell death by inhibiting Trem2 during early atheroma development may slow progression, agonizing Trem2 in advanced lesions may be more important to promote plaque stability.

Mechanistically, Trem2 agonism reprograms foamy macrophage metabolism toward oxidative phosphorylation (OXPHOS), promotes collagen production and decreases inflammatory responses. Interestingly, both foamy macrophages and smooth muscle cells within atherosclerotic plaques have reduced capacity for OXPHOS leading to enhanced cell death, exacerbated necrotic core formation, and decreased fibrous cap size^36^. Similarly, adipose macrophages lacking the OXPHOS pathway undergo excessive ER stress upon lipid challenge due to impaired cholesterol efflux, and subsequent cell death^37^. Moreover, it is well understood that alternatively activated macrophages that promote wound healing switch their metabolism from glycolysis toward oxidative phosphorylation^38^. Thus, we propose that the pro-wound healing response and improved foamy macrophage viability observed with Trem2 agonism may be mediated by shifting metabolism toward OXPHOS. Future studies will need to address this possibility. Furthermore, exactly what molecular mechanisms are driving OXPHOS downstream of Trem2 remain unclear. It has been suggested that the mTOR pathway is induced downstream of Trem2 signaling^22^, which enhances mitochondrial function; however, the link between this pathway and OXPHOS is yet to be established.

Trem2 has been proposed to be a therapeutic target in several diseases, including cancer and Alzheimer’s disease. Trem2 is an attractive target primarily because of its immunomodulatory function under diseased states but non-vital role in acute inflammation such as infection. While we found that AL002a treatment led to expanded plaque size in our murine model, we hypothesize that promoting macrophage survival, decreasing necrotic core formation, and improving ECM production is likely overall beneficial for advanced human atherosclerosis. Importantly, while macrophages are the primary cell subset within murine plaques and early human lesions, in advanced human atheroma’s macrophage content is dramatically less and cellular constituents of plaques are more heterogeneous^39^. Thus, while increasing macrophage cellularity by driving proliferation and survival with Trem2-agonist may lead to plaque expansion in mice, it is likely that this effect in less macrophage dense human plaques would not significantly exacerbate disease burden. However, expanding the pro-wound healing macrophage phenotype downstream of Trem2-agonism to drive collagen production and efferocytosis of dying cells, may indeed be sufficient to provide clinically relevant remodeling to prevent or reduce risk for major cardiovascular events. Overall, this study advances our understanding of how Trem2 signaling vitally regulates foamy macrophage function within atherosclerotic plaques and emphasizes the potential utility of a Trem2 targeting antibody for therapeutic treatment of established atherosclerotic disease.

## Data Availability

Newly generated gene expression data (scRNA-seq) will be made available on the gene expression omnibus (GEO) repository following peer-review of this manuscript. All other data supporting the findings in this study will be made available upon reasonable request to the corresponding author.

## Acknowledgements

We would like to thank Alector, Inc. for supplying Trem2-agonist (AL002a) for this study. This study was funded by grant support from National Institutes of Health (NIH) NIAID R01 AI165553 (JWW), American Heart Association (AHA) CDA855022 (JWW), UMN Dept. of Integrative Biology & Physiology Accelerator Funds, and the Minnesota Office of Higher Education Award (JWW). MTP was supported by a predoctoral research fellowship AHA 903380.

## Author Contributions

JWW, MTP, BB, IT, and DW conceived and designed the research project. MTP, YX, HH, VO, PRS, AEK, AZ, ST, SS, and JWW performed experiments. MTP and JWW wrote the manuscript and prepared figures. All authors assisted with data interpretation and manuscript revision.

## Supplemental Figure Legends

**Supplemental Figure 1:**
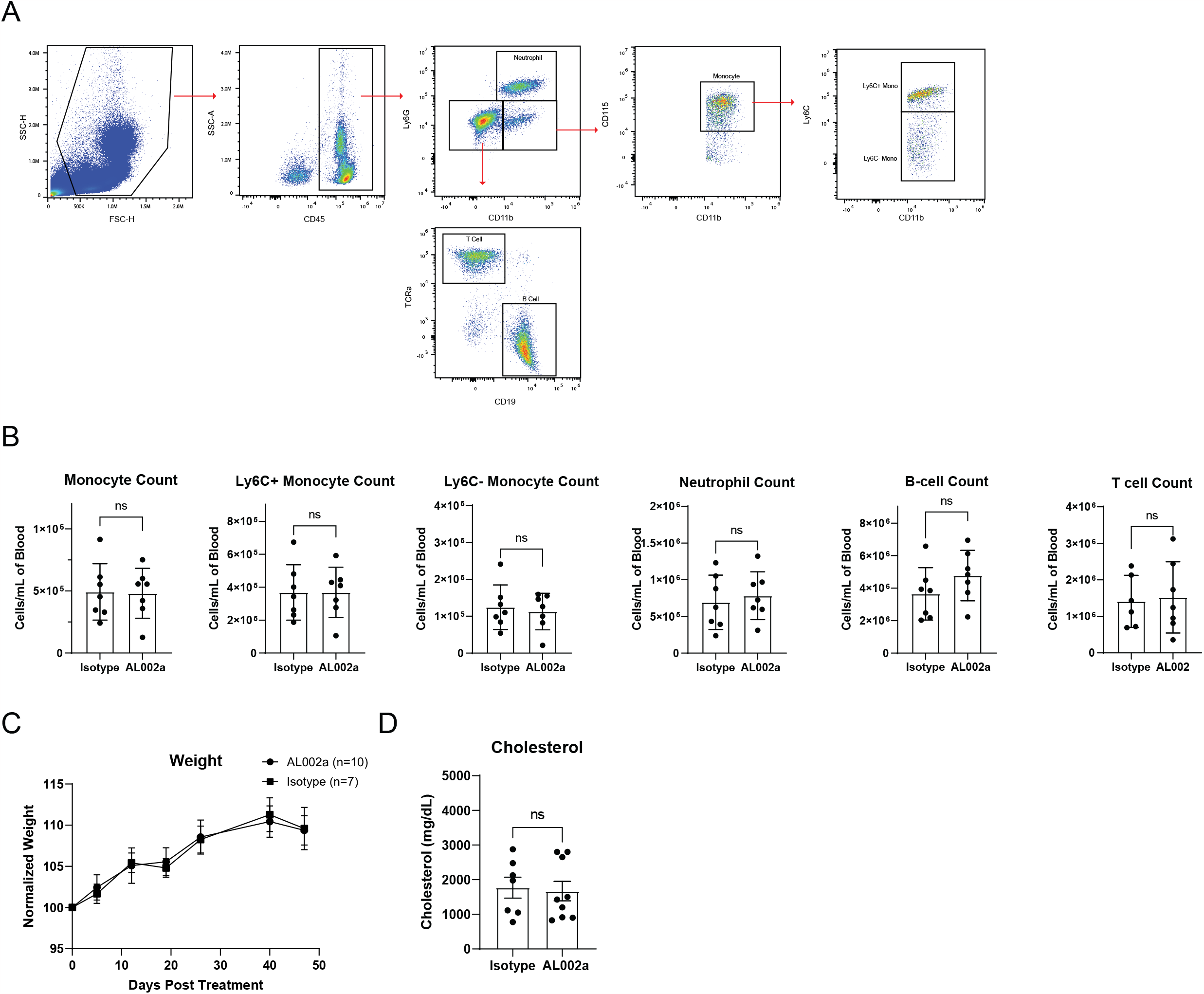
Trem2 agonism does not alter proportions of blood immune cells, body weight or serum cholesterol levels. A) Flow cytometric gating strategy for identifying major blood immune cell populations. B) Blood immune cell profiling by flow cytometry in indicated mice after 16 weeks HFD (n=7/group). Data are mean ± S.E.M. C) Serum cholesterol levels from 16 week HFD fed mice (n=7/group). Data are mean ± S.E.M.

**Supplemental Figure 2:**
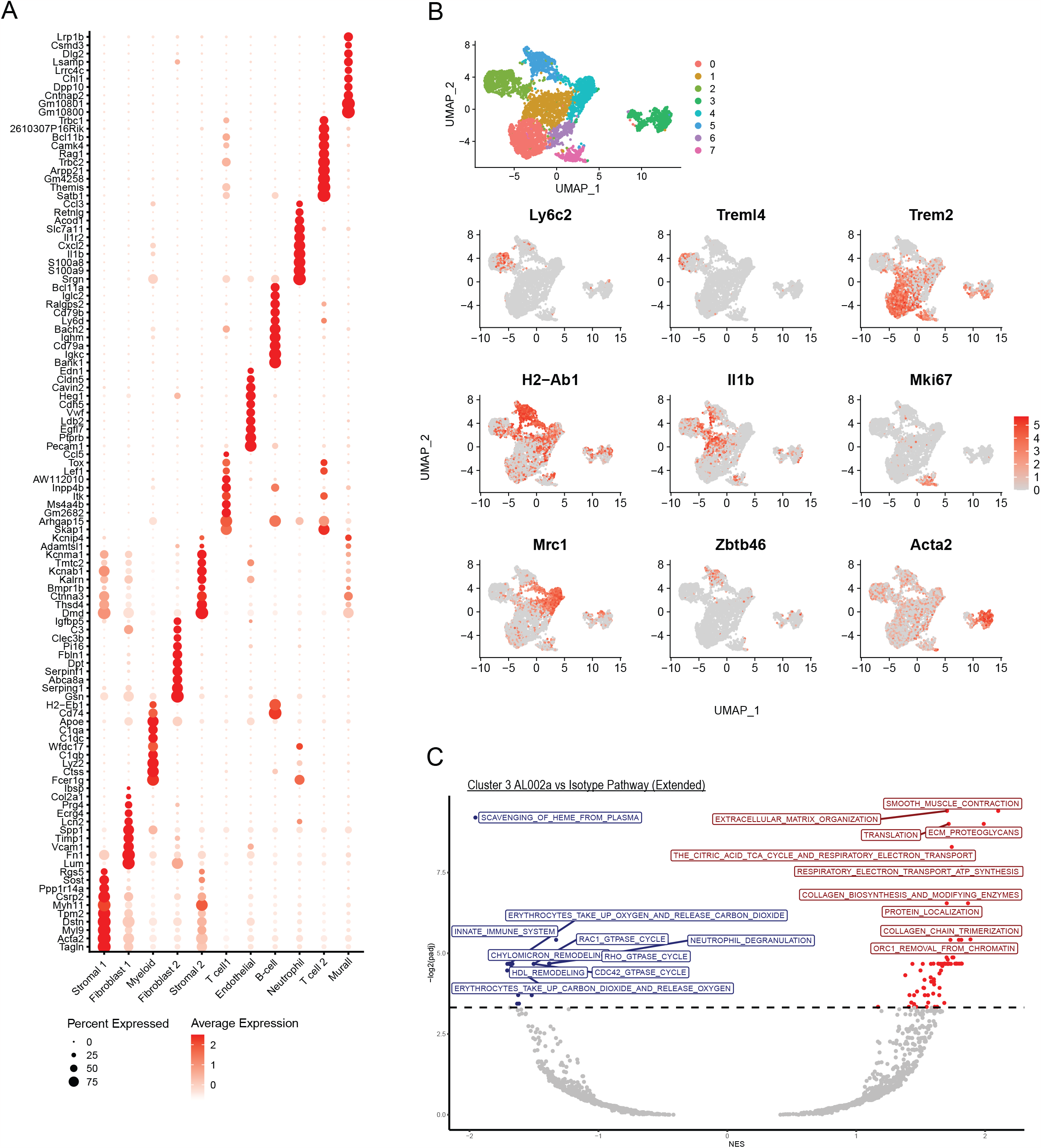
Gene expression from scRNAseq of atherosclerotic aortae. A) Differential gene analysis for each cell cluster from Figure 2B of each main cell population. B) UMAP clustering and gene expression of cluster defining genes from myeloid clustering in Figure 2E. C) Extended REACTOME pathway analysis from Figure 2J showing top 10 upregulated and downregulated pathways between smooth muscle foamy cells from AL002a (red) and Isotype (blue).

**Supplemental Figure 3:**
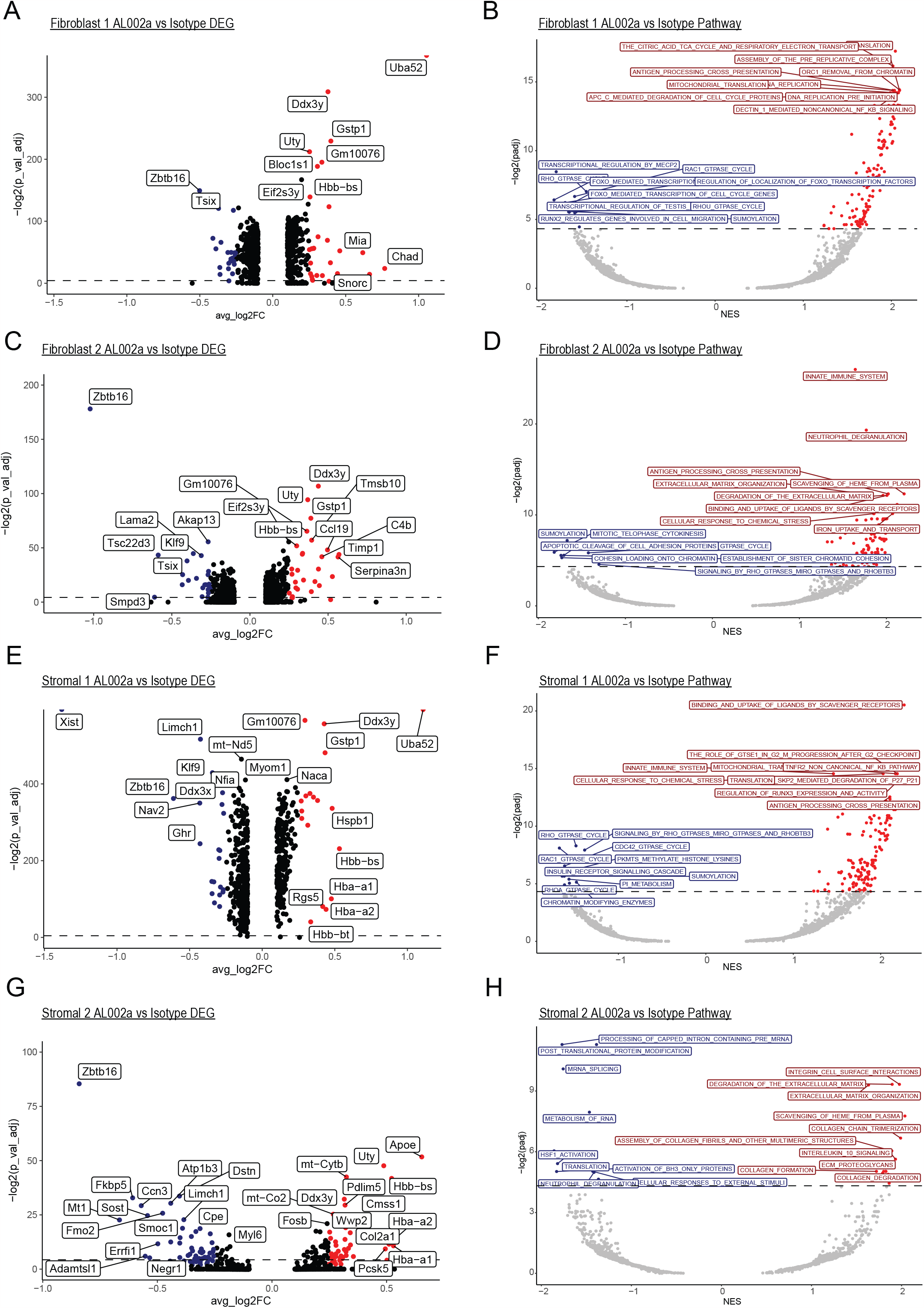
Fibroblast and stromal cells display increased ECM genes with AL002a treatment. A,B) Differential gene expression (A) and REACTOME pathway analysis (B) between fibroblast 1 population of isotype and AL002a treated mice. Red-AL002a, Blue-Isotype. C,D) Differential gene expression (C) and REACTOME pathway analysis (D) between fibroblast 2 population of isotype and AL002a treated mice. Red-AL002a, Blue-Isotype. E,F) Differential gene expression (E) and REACTOME pathway analysis (F) between stromal 1 population of isotype and AL002a treated mice. Red-AL002a, Blue-Isotype. G,H) Differential gene expression (G) and REACTOME pathway analysis (H) between stromal 2 population of isotype and AL002a treated mice. Red-AL002a, Blue-Isotype.

## References

1. Glass CK, Witztum JL. Atherosclerosis. Cell 2001;104:503–516. doi:10.1016/S0092-8674(01)00238-0.

2. Rayner KJ. Cell Death in the Vessel Wall: The Good, the Bad, the Ugly. ATVB 2017;37. doi:10.1161/ATVBAHA.117.309229.

3. Neels JG, Gollentz C, Chinetti G. Macrophage death in atherosclerosis: potential role in calcification. Front Immunol 2023;14:1215612. doi:10.3389/fimmu.2023.1215612.

4. Tabas I. Macrophage death and defective inflammation resolution in atherosclerosis. Nat Rev Immunol 2010;10:36–46. doi:10.1038/nri2675.

5. Tabas I. Consequences and Therapeutic Implications of Macrophage Apoptosis in Atherosclerosis: The Importance of Lesion Stage and Phagocytic Efficiency. ATVB 2005;25:2255–2264. doi:10.1161/01.ATV.0000184783.04864.9f.

6. Seimon T, Tabas I. Mechanisms and consequences of macrophage apoptosis in atherosclerosis. Journal of Lipid Research 2009;50:S382–S387. doi:10.1194/jlr.R800032-JLR200.

7. Thorp E, Subramanian M, Tabas I. The role of macrophages and dendritic cells in the clearance of apoptotic cells in advanced atherosclerosis. Eur J Immunol 2011;41:2515–2518. doi:10.1002/eji.201141719.

8. Boada-Romero E, Martinez J, Heckmann BL, Green DR. The clearance of dead cells by efferocytosis. Nat Rev Mol Cell Biol 2020;21:398–414. doi:10.1038/s41580-020-0232-1.

9. Puylaert P, Zurek M, Rayner KJ, De Meyer GRY, Martinet W. Regulated Necrosis in Atherosclerosis. ATVB 2022;42:1283–1306. doi:10.1161/ATVBAHA.122.318177.

10. Bentzon JF, Otsuka F, Virmani R, Falk E. Mechanisms of Plaque Formation and Rupture. Circ Res 2014;114:1852–1866. doi:10.1161/CIRCRESAHA.114.302721.

11. Lutgens E, Van Suylen R-J, Faber BC, Gijbels MJ, Eurlings PM, Bijnens A-P, et al. Atherosclerotic Plaque Rupture: Local or Systemic Process? ATVB 2003;23:2123–2130. doi:10.1161/01.ATV.0000097783.01596.E2.

12. Ross R. Atherosclerosis — An Inflammatory Disease. N Engl J Med 1999;340:115–126. doi:10.1056/NEJM199901143400207.

13. Falk E, Shah PK, Fuster V. Coronary Plaque Disruption. Circulation 1995;92:657–671. doi:10.1161/01.CIR.92.3.657.

14. Cai B, Thorp EB, Doran AC, Sansbury BE, Daemen MJAP, Dorweiler B, et al. MerTK receptor cleavage promotes plaque necrosis and defective resolution in atherosclerosis. J Clin Invest 2017;127:564–568. doi:10.1172/JCI90520.

15. Jarr K-U, Nakamoto R, Doan BH, Kojima Y, Weissman IL, Advani RH, et al. Effect of CD47 Blockade on Vascular Inflammation. N Engl J Med 2021;384:382–383. doi:10.1056/NEJMc2029834.

16. Kojima Y, Volkmer J-P, McKenna K, Civelek M, Lusis AJ, Miller CL, et al. CD47-blocking antibodies restore phagocytosis and prevent atherosclerosis. Nature 2016;536:86–90. doi:10.1038/nature18935.

17. Endo-Umeda K, Kim E, Thomas DG, Liu W, Dou H, Yalcinkaya M, et al. Myeloid LXR (Liver X Receptor) Deficiency Induces Inflammatory Gene Expression in Foamy Macrophages and Accelerates Atherosclerosis. ATVB 2022;42:719–731. doi:10.1161/ATVBAHA.122.317583.

18. Joseph SB, McKilligin E, Pei L, Watson MA, Collins AR, Laffitte BA, et al. Synthetic LXR ligand inhibits the development of atherosclerosis in mice. Proc Natl Acad Sci USA 2002;99:7604–7609. doi:10.1073/pnas.112059299.

19. Tangirala RK, Bischoff ED, Joseph SB, Wagner BL, Walczak R, Laffitte BA, et al. Identification of macrophage liver X receptors as inhibitors of atherosclerosis. Proc Natl Acad Sci USA 2002;99:11896–11901. doi:10.1073/pnas.182199799.

20. Barrett TJ. Macrophages in Atherosclerosis Regression. ATVB 2020;40:20–33. doi:10.1161/ATVBAHA.119.312802.

21. Zernecke A, Winkels H, Cochain C, Williams JW, Wolf D, Soehnlein O, et al. Meta-Analysis of Leukocyte Diversity in Atherosclerotic Mouse Aortas. Circ Res 2020;127:402–426. doi:10.1161/CIRCRESAHA.120.316903.

22. Ulland TK, Song WM, Huang SC-C, Ulrich JD, Sergushichev A, Beatty WL, et al. TREM2 Maintains Microglial Metabolic Fitness in Alzheimer’s Disease. Cell 2017;170:649–663.e13. doi:10.1016/j.cell.2017.07.023.

23. Hamerman JA, Jarjoura JR, Humphrey MB, Nakamura MC, Seaman WE, Lanier LL. Cutting Edge: Inhibition of TLR and FcR Responses in Macrophages by Triggering Receptor Expressed on Myeloid Cells (TREM)-2 and DAP12. The Journal of Immunology 2006;177:2051–2055. doi:10.4049/jimmunol.177.4.2051.

24. Yao H, Coppola K, Schweig JE, Crawford F, Mullan M, Paris D. Distinct Signaling Pathways Regulate TREM2 Phagocytic and NFκB Antagonistic Activities. Front Cell Neurosci 2019;13:457. doi:10.3389/fncel.2019.00457.

25. Wang S, Mustafa M, Yuede CM, Salazar SV, Kong P, Long H, et al. Anti-human TREM2 induces microglia proliferation and reduces pathology in an Alzheimer’s disease model. Journal of Experimental Medicine 2020;217:e20200785. doi:10.1084/jem.20200785.

26. Patterson MT, Firulyova MM, Xu Y, Bishop C, Zhu A, Schrank PR, et al. Trem2 Promotes Foamy Macrophage Lipid Uptake and Survival in Atherosclerosis.BioRxiv 2022:2022–11.

27. Piollet M, Porsch F, Rizzo G, Kapser F, Schulz DJJ, Kiss MG, et al. TREM2 limits necrotic core formation during atherogenesis by controlling macrophage survival and efferocytosis. Immunology; 2023. doi:10.1101/2023.05.15.539977.

28. Schlegel M, Sharma M, Brown EJ, Newman AAC, Cyr Y, Afonso MS, et al. Silencing Myeloid Netrin-1 Induces Inflammation Resolution and Plaque Regression. Circ Res 2021;129:530–546. doi:10.1161/CIRCRESAHA.121.319313.

29. Kim K, Shim D, Lee JS, Zaitsev K, Williams JW, Kim K-W, et al. Transcriptome Analysis Reveals Nonfoamy Rather Than Foamy Plaque Macrophages Are Proinflammatory in Atherosclerotic Murine Models. Circ Res 2018;123:1127–1142. doi:10.1161/CIRCRESAHA.118.312804.

30. Zhao N, Qiao W, Li F, Ren Y, Zheng J, Martens YA, et al. Elevating microglia TREM2 reduces amyloid seeding and suppresses disease-associated microglia. Journal of Experimental Medicine 2022;219:e20212479. doi:10.1084/jem.20212479.

31. Aran D, Looney AP, Liu L, Wu E, Fong V, Hsu A, et al. Reference-based analysis of lung single-cell sequencing reveals a transitional profibrotic macrophage. Nat Immunol 2019;20:163–172. doi:10.1038/s41590-018-0276-y.

32. Newman AAC, Serbulea V, Baylis RA, Shankman LS, Bradley X, Alencar GF, et al. Multiple cell types contribute to the atherosclerotic lesion fibrous cap by PDGFRβ and bioenergetic mechanisms. Nat Metab 2021;3:166–181. doi:10.1038/s42255-020-00338-8.

33. Nugent AA, Lin K, Van Lengerich B, Lianoglou S, Przybyla L, Davis SS, et al. TREM2 Regulates Microglial Cholesterol Metabolism upon Chronic Phagocytic Challenge. Neuron 2020;105:837–854.e9. doi:10.1016/j.neuron.2019.12.007.

34. Kockx M. Apoptosis in atherosclerosis: beneficial or detrimental? Cardiovascular Research 2000;45:736–746. doi:10.1016/S0008-6363(99)00235-7.

35. Geng Y-J, Libby P. Progression of Atheroma: A Struggle Between Death and Procreation. ATVB 2002;22:1370–1380. doi:10.1161/01.ATV.0000031341.84618.A4.

36. Yu EPK, Reinhold J, Yu H, Starks L, Uryga AK, Foote K, et al. Mitochondrial Respiration Is Reduced in Atherosclerosis, Promoting Necrotic Core Formation and Reducing Relative Fibrous Cap Thickness. ATVB 2017;37:2322–2332. doi:10.1161/ATVBAHA.117.310042.

37. Wculek SK, Heras-Murillo I, Mastrangelo A, Mañanes D, Galán M, Miguel V, et al. Oxidative phosphorylation selectively orchestrates tissue macrophage homeostasis. Immunity 2023;56:516–530.e9. doi:10.1016/j.immuni.2023.01.011.

38. Viola A, Munari F, Sánchez-Rodríguez R, Scolaro T, Castegna A. The Metabolic Signature of Macrophage Responses. Front Immunol 2019;10:1462. doi:10.3389/fimmu.2019.01462.

39. Fernandez DM, Rahman AH, Fernandez NF, Chudnovskiy A, Amir ED, Amadori L, et al. Single-cell immune landscape of human atherosclerotic plaques. Nat Med 2019;25:1576–1588. doi:10.1038/s41591-019-0590-4.

